# Analysis of the potential of environmental *Burkholderia sensu lato* isolates to cause infection in a zebrafish embryo model

**DOI:** 10.1101/2024.12.26.630235

**Authors:** J Gigan, J Boyer, L Moulin, AC Vergunst

## Abstract

*Burkholderia sensu lato* (s.l.) regroups several genera of closely related bacterial species with remarkable diversity in metabolic features. They can be free-living or have mutualistic or pathogenic interactions with different host organisms, including insects, plants and animals. *Burkholderia* s.l. species display numerous characteristics that can be beneficial for agronomical and biotechnological applications, including promotion of plant growth and bioremediation; however, many of the species are known human pathogens. Several recent taxonomical reclassifications of the *Burkholderia* genus resulted in its separation into 7 distinct genera. *Burkholderia sensu stricto* (s.s.), which includes the *Burkholderia cepacia* complex and the *pseudomallei* group, regroups most of the species that have been shown to cause infection in humans and plants. Little is known about the pathogenic potential of species belonging to the other 6 genera; however there have been occasional reports of human infections.

Here, the vertebrate zebrafish embryo was used to analyse the behaviour of a panel of environmental isolates belonging to different species of the *Burkholderia cepacia* complex (5), *Paraburkholderia* (5) and *Caballeronia* (1). *Burkholderia cepacia* P14-NS, *Burkholderia orbicola* P21-NS, and *Paraburkholderia* sp. ABIP659 (closest to *P. tropica*, and further named *tropica*-like) each caused a low percentage (2%) of embryo mortality. While the other strains had not killed the embryos during the 4-day time span of the experiments, overall, *Paraburkholderia* and *Burkholderia* s.s. strains persisted equally well in this vertebrate host. Unexpectedly, *P. tropica*-like showed the highest increase in bacterial burden during infection, and induced the strongest pro-inflammatory response, including abscess formation, and poor prognosis at the endpoint of the experiment. Our study emphasizes that in vivo infection studies are needed to gain more insight into potential pathogenicity of the different *Burkholderia* s.l. species.

## Introduction

*Burkholderia* s.l. is composed of a large group of Gram-negative β-proteobacteria that are ubiquitously found in nature, often in the rhizosphere of plants including rice, maize and onions, but also isolated as symbionts or pathogens from other ecological niches, such as plants, insects, fungi and opportunistic human infections. *Burkholderia cepacia*, initially named *Pseudomonas cepacia*, was first described as a plant pathogen causing sour skin in onion ^1^. This group of highly versatile bacteria includes species displaying valuable biotechnological and agricultural properties such as plant growth promotion and antifungal activity, but also species that are pathogens of plants and animals. Some *Burkholderia* strains were used in the 90s for their biocontrol capacity against several crop pathogens, however, they were withdrawn from the market by the United States Environmental Protection Agency (EPA) after risk assessment due to the presence of major pathogens for cystic fibrosis (CF) patients and immunocompromised individuals in the *Burkholderia* genus (https://www.gpo.gov/fdsys/pkg/FR-2004-09-29/pdf/04-21695.pdf).

Over the past 10 years, in face of the dilemma of species with large agricultural, industrial and biotechnological potential versus the presence of species that are major human pathogens, the *Burkholderia* s.l. has seen several taxonomic reclassifications. The most recent taxonomic changes were based on whole genome sequences and phenotypic characteristics, and finally resulted in the division of *Burkholderia* s.l. into 7 different genera: *Burkholderia sensu stricto* (s.s.), *Paraburkholderia, Caballeronia, Robbsia*, *Pararobbsia*, *Trinickia* and *Mycetohabitans* ^2–6^, with currently more than 4000 associated genome assemblies ^7^.

The distinction of species as either pathogenic or environmental-beneficial based on their phylogenetic classification (*Burkholderia* s.s. versus the other 6 *Burkholderia* s.l. species, respectively), has been controversial ^8,9^. It is often considered that species from the “environmental-beneficial group” do not pose a health hazard for humans and are safe for agricultural and industrial applications ^2,10–12^. However, although infrequently, *Paraburkholderia* and *Caballeronia* strains have been isolated from human opportunistic infections; *P. tropica*, described as a plant-growth promoting bacteria for barley ^13^, has been isolated from a human infection ^14^ as have *P. fungorum* ^15,16^, *C. turbans* and *C. concitans* ^17^. A separation into pathogenic and environmental-beneficial groups can be further misleading since *Burkholderia* s.s. species with clinical importance are environmental bacteria of which many have been shown to have beneficial properties^,8,18,19^.

The majority of human pathogens, including the *Burkholderia cepacia* complex (Bcc), belong to the genus *Burkholderia* s.s., which is currently composed of 26 closely related species. The Bcc are important pathogens of CF patients, especially *B. cenocepacia* and *B. multivorans*, and *Burkholderia gladioli*, a non-Bcc member ^20^. High levels of transmissibility from patient to patient and their intrinsic resistance to commonly used antibiotics which complicates treatment of these infections are correlated with poor clinical prognosis ^21^. In addition, patients with chronic granulomatous disease and immunocompromised individuals are also vulnerable to infections with Bcc ^22^. *B. cepacia* is a major contaminant of pharmaceutical industry and increasing cases of nosocomial and community acquired infections are being reported ^23,24^. Community acquired Bcc infections are an important factor of concern, mainly through contaminated pharmaceuticals, and cases of Bcc infection in immune-competent individuals are also reported ^25–27^.

*B. cenocepacia* can persist and create an intracellular replication niche in phagocytic and non-phagocytic cells (reviewed in ^28^). In vivo infection models, including *Drosophila*, *Caenorhabditis elegans*, *Galleria mellonella*, plants, rodents, and *Danio rerio* (zebrafish) ^29–37^, have provided insight into bacterial virulence determinants and host-pathogen interactions in the context of a whole organism. However, infection studies of *Burkholderia* s.l. species of the “environmental” group are limited to studies using amoebae and nematodes ^5,10,38^.

The zebrafish is an established vertebrate model for infection studies, particularly to study the interaction of microorganisms with the host innate immune system using non-invasive real time imaging using the optically transparent embryos ^39,40^. Significant homology between the zebrafish inflammatory and immune signalling pathways with those of humans have resulted in better understanding of cellular and molecular basis of infection mechanisms ^41–43^. We reported earlier that zebrafish embryos have helped demonstrate that clinical isolates of the ET-12 lineage, including *B. cenocepacia* K56-2 and J2315, are able to cause a rapidly fatal acute pro-inflammatory infection, while other clinical isolates, including *B. stabilis* LMG14294 and *B. vietnamiensis* FC441, cause persistent infection with a constant bacterial burden without any obvious inflammatory responses ^34,44^. An intramacrophage replication stage was essential for the development of pro-inflammatory fatal infection by *B. cenocepacia* in zebrafish embryos ^44^. Importantly, the intracellular nature of Bcc bacteria within alveolar macrophages has been shown in lung tissue samples from CF patients ^45,46^.

In this study, a panel of environmental strains belonging to the genera *Paraburkholderia* (5), *Caballeronia* (1) and the *Burkholderia cepacia* complex (5) have been analysed for virulence potential in the zebrafish embryo model. Host survival and bacterial burden, in vivo observations, neutrophil quantification and host inflammatory gene expression assays showed that most *Burkholderia* and *Paraburkholderia* strains were equally able to survive and persist in the zebrafish embryo. The data show that a *P. tropica-*like strain was the most virulent strain of the analysed panel, causing pro-inflammatory responses leading to abscess formation and poor survival prognosis in zebrafish embryos.

## Materials and Methods

### Ethics statement

The zebrafish facility in the laboratory was approved by the “Direction Départementale de la Protection des Populations” (DDPP) du Gard (ID: 30-189-4). Zebrafish were kept and handled in compliance with the guidelines of the European Union for handling laboratory animals (Animal Protection Directive 2010/63/EU). Infection experiments using zebrafish embryos were terminated 5 days post fertilization and did not require authorization by the ethical committee.

### Zebrafish maintenance and care

Adult zebrafish were maintained with a 14/10 h Light/dark cycle and standard water parameters 28°C, pH 7.3, conductivity 450µS. AB fish were purchased from the Zebrafish International Resource Center (ZIRC). Transgenic reporter lines Tg(*mpx:eGFP*^i114^) ^47^ and Tg(*mpeg1:mCherryF*)^ump2Tg 48^ were used to study interactions with neutrophils and macrophages, respectively. Eggs were obtained by natural spawning and kept in petri dishes containing E3 medium (5mM NaCl, 0.17mM KCl, 0.33mM CaCl2, 0.33mM MgSO4, 10 mM HEPES, pH=7.2) in an incubator at 28°C and with 14/8 h light/dark cycles using a combination of T5 and T8 LEDs. Embryos were anesthetized in E3 medium containing 0.02% buffered MS222 (Ethyl 3-aminobenzoate methane sulfonate (Sigma) during bacterial injections, imaging and fixation.

### Bacterial strains

The strains used in this study are listed in **Table 1**. The optimal medium composition for each strain is indicated in the Table. *B. cenocepacia* K56-2 was grown at 37 °C, all other strains were grown at 30 °C on LB Miller: 10 g/L peptone, 5 g/L yeast extract, 10g/L NaCl +1.5% agar. Some strains were grown in low salt (10g/L NaCl) LB. Tryptic Soy medium (TS) contains per L: 15 g casein, 5 g peptic digest of soybean meal, 5g NaCl, and 1.5% agar. All strains were transformed with plasmid pIN29 ^34^, which encodes the *DSRed* reporter gene. The medium was supplemented with 30 mg/L chloramphenicol (Cm) for the growth of pIN29-derivative strains. For infection experiments, bacterial strains were first grown on solid agar plates from −80 °C glycerol stocks using the appropriate medium for each strain (**Table 1**), followed by overnight growth in liquid LB Miller, except for strain *C. insecticola*, for which modified LB was used. Live dead staining (BacLight Invitrogen) showed that more than 90% of bacteria were alive for all strains after overnight growth in LB liquid (data not shown).

**Table 1.** Strains used in this study. ^a^ Original isolate and ABIP collection numbers and ^b^ ABIP pIN29-derivatives were acquired from the PHIM ABIP collection. RPE75 was obtained from P. Mergaert. ^c^ Culture conditions used for infection experiments as detailed in M&M. LBm = modified LB. CF=cystic fibrosis.

| *Burkholderia* s.l. species | Strain <sup>a</sup> | Source | Dsred (pIN29) derivative <sup>b</sup> | Culture medium <sup>c</sup> | References |
| --- | --- | --- | --- | --- | --- |
| *Paraburkholderia caribensis* | ABIP 1334 | *Oryza sativa* | ABIP 2320 | TSA 50% | <sup>50</sup> |
| *Paraburkholderia graminis* | C4D1M<sup>T</sup> = ABIP 5 | Maize root | ABIP 560 | LBm | <sup>51</sup> |
| *Paraburkholderia kururiensis* | M130 = ABIP 1108 | *Oryza sativa* endophyte | ABIP 2202 | LBm | <sup>52</sup> |
| *Paraburkholderia phytofirmans* | PsJN<sup>T</sup> = ABIP 23 | Plant Endophyte | ABIP 527 | LBm | <sup>50,53</sup> |
| *Paraburkholderia* sp, (closest to *P. tropica*), named *tropica*-like | ABIP 659 | *Oryza sativa* rhizosphere | ABIP 1121 | modified LB | <sup>54</sup> |
| *Caballeronia insecticola* | RPE64<sup>T</sup> = ABIP2336 | *Riptortus pedestris* | RPE75 | modified LB | <sup>55</sup> |
| *Burkholderia cepacia* | ABIP 441 | *Oryza sativa* rhizosphere | ABIP 2318 | modified LB | <sup>54</sup> |
| *Burkholderia orbicola* | ABIP 444 | *Oryza sativa* root | ABIP 2321 | LB Miller | <sup>56</sup> |
| *Burkholderia vietnamiensis* | LMG 10929<sup>T</sup> = ABIP 47 | *Oryza sativa*, rhizosphere soil | ABIP 572 | LB Miller | <sup>57</sup> |
| *B. sp. 3* (closest to *multivorans*) | ABIP 3274 | *Oryza sativa* root | ABIP 3292 | TSA 50% | L. Moulin |
| *B. sp. 2* (closest to *multivorans*) | ABIP 3277 | *Oryza sativa* root | ABIP 3298 | TSA 50% | L. Moulin |
| *B. cenocepacia* | K56-2 LMG 18863 | CF isolate | K56-2 (pIN29) | LB Miller | <sup>58</sup> |
| *B. stabilis* | LMG14294 | CF isolate | LMG12494 (pIN29) | LB Miller | <sup>59</sup> |

### Zebrafish infection assays

Zebrafish eggs were dechorionated manually with fine tweezers (Dumont no 5) 2 hours prior to infection. Embryos, staged 28-30 hpf, were arranged in 1 mm slots on 1.5% agarose E3 plates prepared using molds (IBI Scientific) and supplemented with E3 medium containing 0.02% buffered MS222, and injected intravenously with 200-400 CFU in PBS containing 0.1% phenol red as described ^49^. To determine the precise inoculum for each strain, injections were performed directly on LB agar plates (n=3; indicated in the legends to the figures). Embryos were incubated in 48 well plates in an incubator at 28°C and with 14/8 h light/dark cycles. Mortality was monitored by confirming the absence of a heartbeat in infected embryos using a binocular microscope. Bacterial burden was determined at 1, 2, 3, and 4 days post fertilization (dpf) by plating embryos manually disrupted in sterile H_2_O using Eppendorf Pestles (5 individual embryos per time point). For each strain the appropriate media were used for optimal recovery of bacteria from embryos (Table 1).

### RNA extraction, cDNA synthesis and quantitative RT-PCR

RNA extraction, cDNA synthesis and qRT-PCR analysis were performed as described previously ^49^. Three biological replicates were performed, each with three technical replicates per sample, unless otherwise mentioned. As housekeeping gene the peptidylprolyl isomerase A-like (*ppial*) gene was used. The ΔΔCt method was used for analysis of the data, and log_2_-transformed fold changes are represented as column bar graphs normalized to a PBS-injected control group at each time point.

### Microscopy imaging

A Leica DM IRB inverted microscope with a Coolsnap fx camera (Roper Scientific) and a Nikon AZ100, coupled with a Coolsnap HQ2 (Roper Scientifique) were used to image embryos using MetaVue software. Images at different time points show independent embryos. Figures were prepared using Adobe Photoshop. A confocal microscope (FV10i with Fluoview, Olympus) and Imaris (Andor Technology Ltd) were used to acquire image 5B.

### Quantification of neutrophils

For quantification of neutrophil numbers, Tg(mpx:eGFP^i114^) embryos were injected with bacterial strains. At different time points after infection, on average 10 embryos per time point per strain were imaged using an AZ100 fluorescence microscope, using similar settings for images at each time point. The images were analysed in Imaris using the module “spots”, to count individual neutrophils.

### Statistical analysis

GraphPad Prism 10 software was used for statistical analysis. Survival was analysed using a Log rank (Mantel-Cox) test. Bacterial burden in individual embryos is represented in dot plots (Figure 1). The average inoculum at T=0 is calculated from 3 injections on agar plate, and indicated for each strain in the legend. CFU counts from individual embryos at later time points were log_10_ transformed and indicated as individual dots. One-way ANOVA with Sidak’s Multiple Comparisons test was used to determine any differences in bacterial burden at day 1, and day 3 for each strain. To include embryos in which no CFU were detected, the 0 count was converted to 1, prior to log-transformation. Neutrophil cell counts were analysed using one-way ANOVA with Sidak’s Multiple Comparisons test. Significance of the qRT-PCR data was analysed using one-way ANOVA with Tukey’s Multiple Comparison Test. Columns indicate mean fold with SEM. For each treatment, normalized to the corresponding PBS control, significance in relative fold-change is indicated with an asterisk above the column. Significance is indicated as: ns, non-significant, *, p ≤ 0.05; **, p ≤ 0.01; ***, p ≤ 0.001; ****, p ≤ 0.0001.

**Figure 1.**
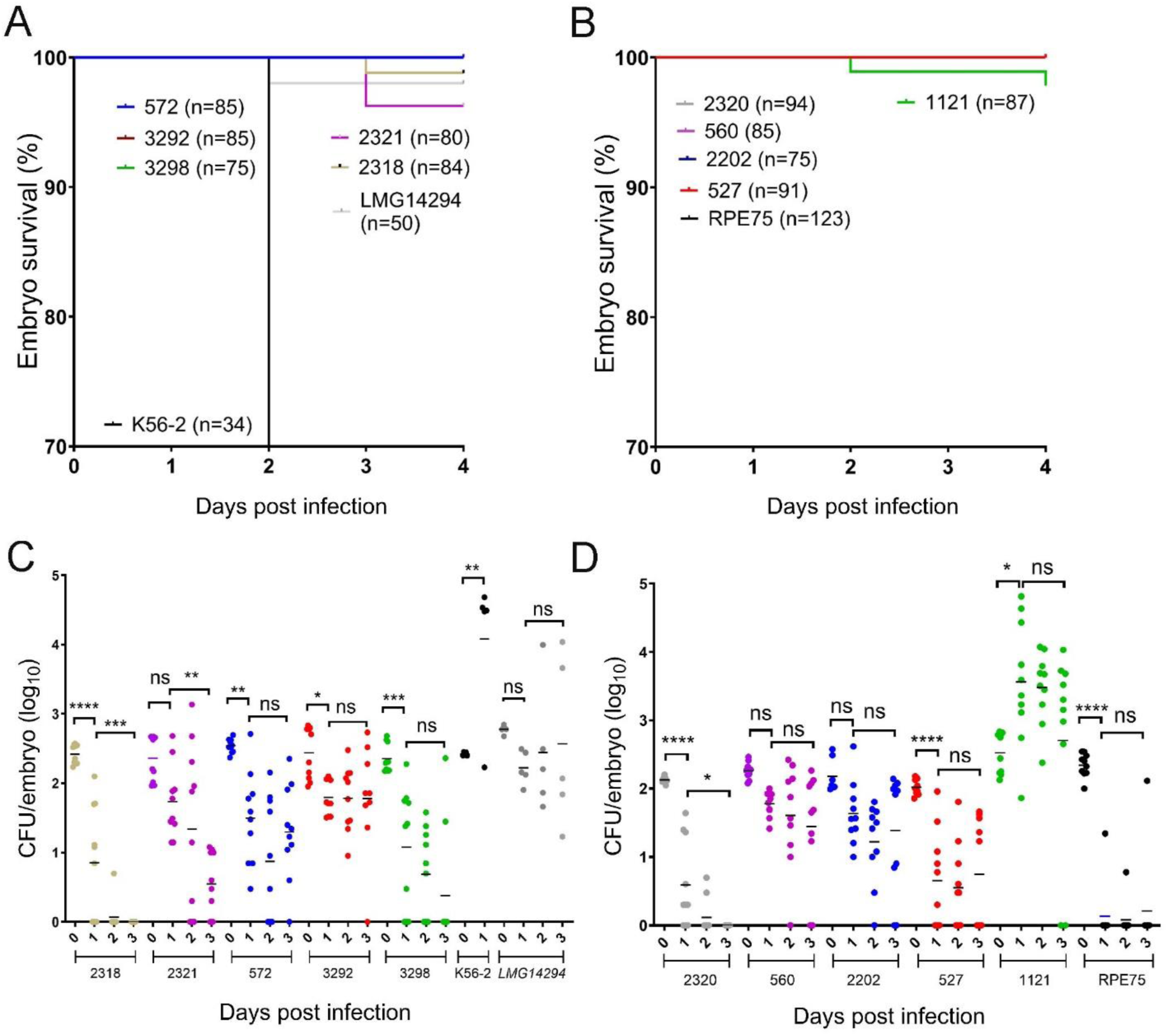
Embryo survival after injection with **A.** *Burkholderia* strains and **B.** *Paraburkholderia* and *Caballeronia strains* (see Table 1, ABIP numbers), pooled data from 2 experiments, n= number of individual embryos analysed. The Y-axe is cut off at 70% to better visualise the low levels of mortality. **C, D**, Dot plots showing bacterial burden (same experiments as shown in A and B, respectively) over 3 days. **C**, *Burkholderia strains* (average inoculum: ABIP 2318: 275 CFU; ABIP 2321: 287 CFU; ABIP 572: 355 CFU; ABIP 3292: 368 CFU; ABIP 3298: 252 CFU; *B. cenocepacia* K56-2: 265 CFU; *B. stabilis* LMG12494: 605 CFU). D, *Paraburkholderia* and *Caballeronia* strains (average inoculum: ABIP 2320: 135 CFU; ABIP 560: 190 CFU; ABIP 2202: 177 CFU; ABIP 527: 108 CFU; ABIP 1121: 405 CFU; RPE75: 235 CFU). Each data point represents an individual embryo (n=5 per time point per strain per experiment), except T=0 (inoculum directly injected on agar plates. Strains *B. cenocepacia* K56-2, causing rapidly fatal infection, and *B. stabilis* LMG14294, causing persistent infection ^34^, were included in the experiments for direct comparison only (1 experiment). One-way ANOVA per group, significance is indicated between day 0 and 1, and day 1 and 3 resp. * p ≤ 0.05, ** p ≤ 0.01; *** p ≤ 0.001; **** p ≤ 0.0001; ns: not significant.

## Results

### *Paraburkholderia*, *Caballeronia* and Bcc strains persist in zebrafish embryos

Here, we studied the virulence of a panel of 11 environmental *Burkholderia* s.l. isolates (**Table 1**) in a zebrafish embryo model. The panel included 5 environmental Bcc strains isolated from rice (*Oryza sativa*) roots or rhizosphere from different geographic locations: *B. vietnamiensis* (LMG 10929) was isolated in Vietnam, *B. cepacia* (ABIP 441) and *B. orbicola* ABIP 444 were isolated in Cameroon, and 2 strains closest to *B. multivorans* (ABIP 3274, ABIP 3277) were isolated in Cambodia. *B. vietnamiensis* and *B. cenocepacia* are known to cause tissue water-soaking in onion, but they have also been isolated from healthy plants and from human and animal infections. Three *Paraburkholderia* strains, *P. kururiensis* M130, a rice endophyte, *P. caribensis* ABIP 1334 and *Paraburkholderia sp.* ABIP 659 (closest to *tropica* and here named *tropica*-like) were isolated from rice plants in Brazil, Burkina Faso and Vietnam respectively. *P. graminis* C4D1M^T^ was isolated from maize roots in France and *P. phytofirmans* PsJN from onion roots*. C. insecticola* RPE64 is a symbiont needed for normal development of the stink bug *Riptortus pedestris*. We also included two clinical Bcc isolates that we have studied in the zebrafish embryo model in detail; *B. cenocepacia* K56-2, which causes rapidly fatal infection, and *B. stabilis* LMG12494, which leads to persistent infections with constant bacterial load and low pro-inflammatory responses ^34,44^. All strains were electroporated with plasmid pIN29, harbouring a DsRed reporter gene ^34^ (Table 1), and the DsRed-expressing derivative strains were used for the experiments described below.

Embryos, staged at 28-30 hpf, were micro-injected intravenously with the pIN29-containing strains (Table 1) and analysed for survival, bacterial burden and using real time imaging. Analysis of host survival (**Figure 1A and 1B**) showed that 8 of the tested strains did not induce mortality in embryos up to 4 days post infection (dpi), while two *Burkholderia* strains, *B. cepacia* ABIP 2318 and *B. orbicola* ABIP 2321, and one *Paraburkholderia* strain, *P. tropica*-like ABIP 1121, gave low mortality of 2-4 %. Fluorescence imaging showed that death of the embryos infected with *B. cepacia* ABIP 2318 and *B. orbicola* ABIP 2321 was caused by infection and substantial multiplication of the bacteria in the central nervous system (CNS)(**Figure 2**). None of the 11 strains induced 100 % mortality seen with the epidemic isolate *B. cenocepacia* K56-2 (^34^ and **Figure 1A**).

**Figure 2.**
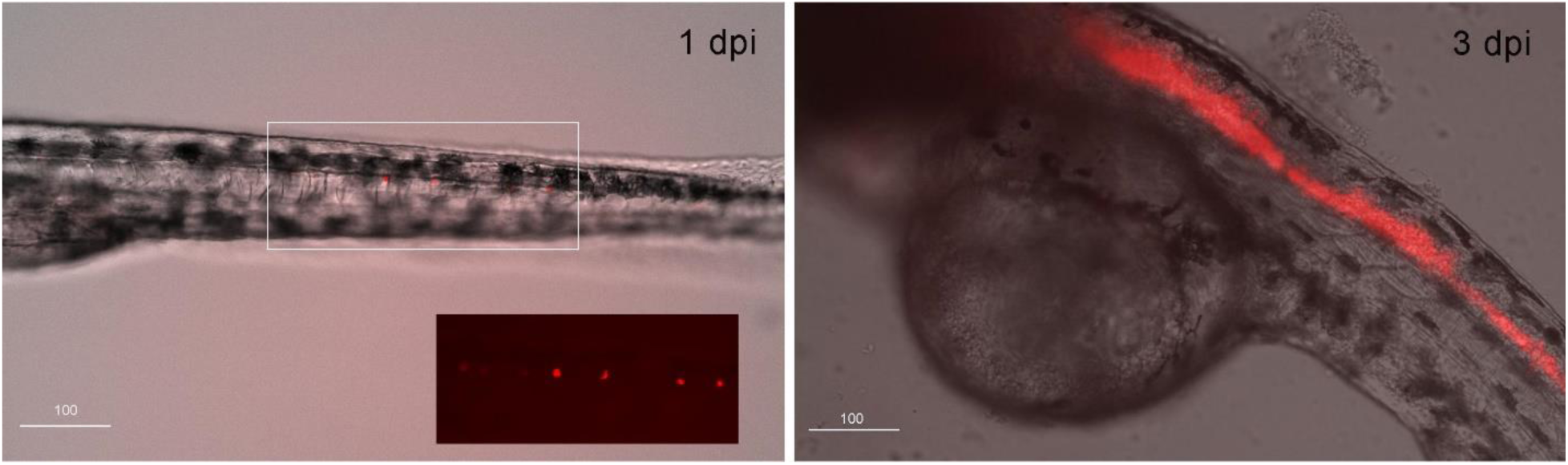
Two zebrafish embryos injected with *B. orbicola* ABIP 2321 expressing DSRed. **A.** Image of embryo caudal region at 1 dpi overlay. Overlay of bright field and fluorescence image, inset shows enlarged image with bacterial clusters, **B.** Embryo at 3 dpi showing infection of the CNS. Scale bars 100 µm.

*B. cepacia* ABIP 2318, despite killing an occasional embryo, and *P. caribensis* ABIP 2320 were completely cleared by infected embryos in 2 to 3 days; however, the other isolates persisted in all or a percentage of zebrafish embryos during the full experimental time course (**Figure 1C and 1D**). Although the bug symbiont *C. insecticola* was also cleared from most infected embryos, occasionally an embryo with the same number of bacteria as the initial inoculum was observed up to 4 days (**Figure 1D** and **3A**). Real time fluorescence imaging at 3 and 4 dpi further showed that persisting bacteria were mostly present in one or multiple micro-colonies, as shown for *C. insecticola* and *B. vietnamiensis* (**Figure 3)**, most likely in macrophages as described below for *P. tropica-*like (**Figure 4B**). Based on the localization of bacteria in several micro clusters at later time points, the constant bacterial burden during infection is likely due to an equilibrium between bacterial killing and replication at low rates.

**Figure 3.**
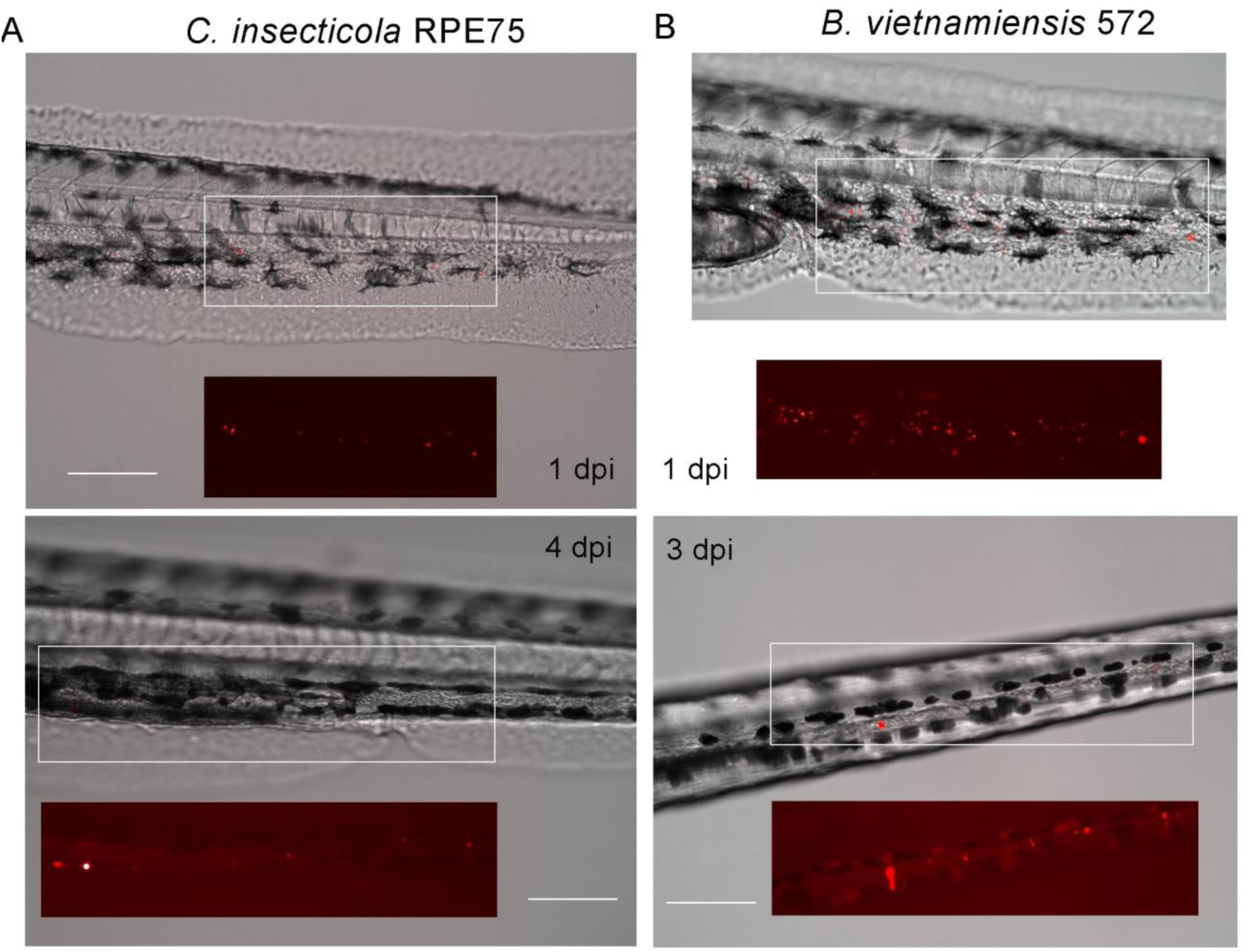
Zebrafish embryos injected with **A.** DSRed expressing *B. insecticola* RPE75, shown at 1 and 4 dpi and **B.** *B. vietnamiensis* ABIP 572. Overlay of bright field and fluorescence images, insets: enlargement of the boxed areas showing the fluorescence image only. Scale bars 100 µm.

**Figure 4.**
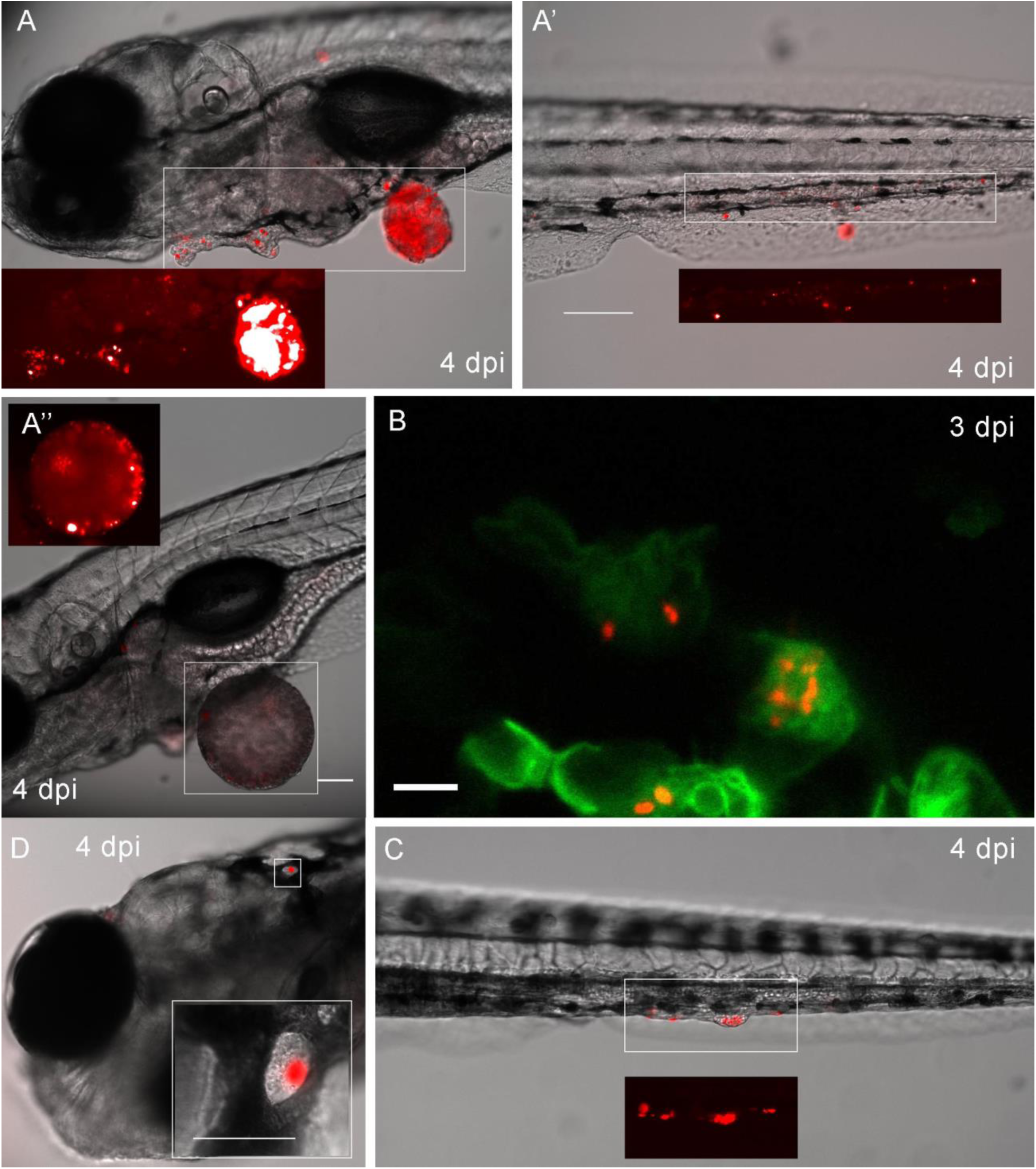
Microscopical observation of DSRed-expressing *P. tropica*-like ABIP 1121 in infected embryos at 4 dpi (**A, A’** and **A’’**). **B.** Close up of infected macrophages in a *mpeg-eGFP* embryo infected with DSRed-expressing *P. tropica* like at 3 dpi. **C.** Zebrafish embryo at 4 dpi infected with DSRed-expressing *P. phytofirmans* ABIP 527. **D.** Embryo infected with *P. kururiensis* ABIP 2202 at 4 dpi. Overlay of bright field and fluorescence images, insets: enlargement of the boxed areas showing the fluorescence images (enhanced signal). Scale bars 100 µm, in B 5 µm. Confocal image (B) shows one Z- image of a confocal stack (green and red filters).

The bacterial burden in embryos infected with *B. vietnamiensis* ABIP 572, *B. multivorans*-like ABIP 3292, *P. graminis* ABIP 560 and *P. kururiensis* ABIP 2202 was stable over 4 days **(Figure 1C and 1D)**. However, *P. tropica*-like ABIP 1121 showed a significant increase in bacterial burden (**Figure 1D**). Although most embryos infected with this strain were still alive at the endpoint of the experiment, microscopical observation showed an uncontrolled bacterial burden in all embryos, and tissue damage and large abscesses filled with bacteria in about 10% of embryos (**Figure 4A**). Although not quantified at this time, confocal imaging showed that the bacteria were present in macrophages (**Figure 4B**). *P. tropica*-like infected embryos showed reduced movement and a poor prognosis at the end of the experiment (not shown).

*P. phytofirmans* ABIP 527 also showed larger infection sites at 4 days post infection in about 10% of the infected embryos (**Figure 4C**). *P. kururiensis* ABIP 2202, an endophyte, was still present in most infected embryos, and observed in clusters at 4 dpi with evidence of replication (**Figure 4D**). Interestingly, two strains, *P. graminis* ABIP 560 and *P. caribensis* ABIP 2320, showed partial resistance to phagocytosis, since free bacteria were still observed moving in the blood circulation after 1 day while bacteria of the other strains were taken up by immune cells (not shown), similar to *B. cenocepacia* K56-2 ^60^.

### Analysis of host pro-inflammatory cytokine gene expression during infection

The pro-inflammatory interleukin Il1b and the chemokine Cxcl8 play critical roles in host immune response to infection and injury and promote neutrophil recruitment to the sites of infection and inflammation. To gain insight in the height of induction of host immune responses to each of the strains, the expression level of *il1b* and *cxcl8* in infected embryos at 3 and 24 hpi was compared to that of PBS-injected embryos (**Figure 5**). Injection of all analysed strains caused significantly increased expression of *il1b* and *cxcl8* at 3 hpi, except *P. kururiensis* ABIP 2202 (**Figure 5)**. As previously reported ^44^, *cxcl8* expression levels in embryos infected with highly virulent *B. cenocepacia* K56-2 remained high at 24 hpi, however, in embryos infected with the other strains, including *B. stabilis* LMG14294, *cxcl8* expression levels had dropped and were not significantly different from PBS-injected controls. As found for *B. stabilis* LMG14294 (**Figure 5A** and ^44^), *il1b* expression at 24 hpi was lower than at 3h. However, unlike *B. stabilis* where the expression level of these two cytokine genes returned to levels of non-infected control embryos, for most *Burkholderia* and *Paraburkholderia* strains of the panel the level remained elevated, with on average 1-2 log_2_-fold higher expression than in PBS-injected controls. Overall, gene expression changes induced by the panel of strains did not differ between *Burkholderia* and *Paraburkholderia*. The highest relative fold-increase for *il1b* expression at 24 hpi was observed in embryos infected with *P. tropica*-like ABIP 1121, in agreement with high bacterial burden (Figure 1B) and microscopical observations (**Figure 4A**).

**Figure 5.**
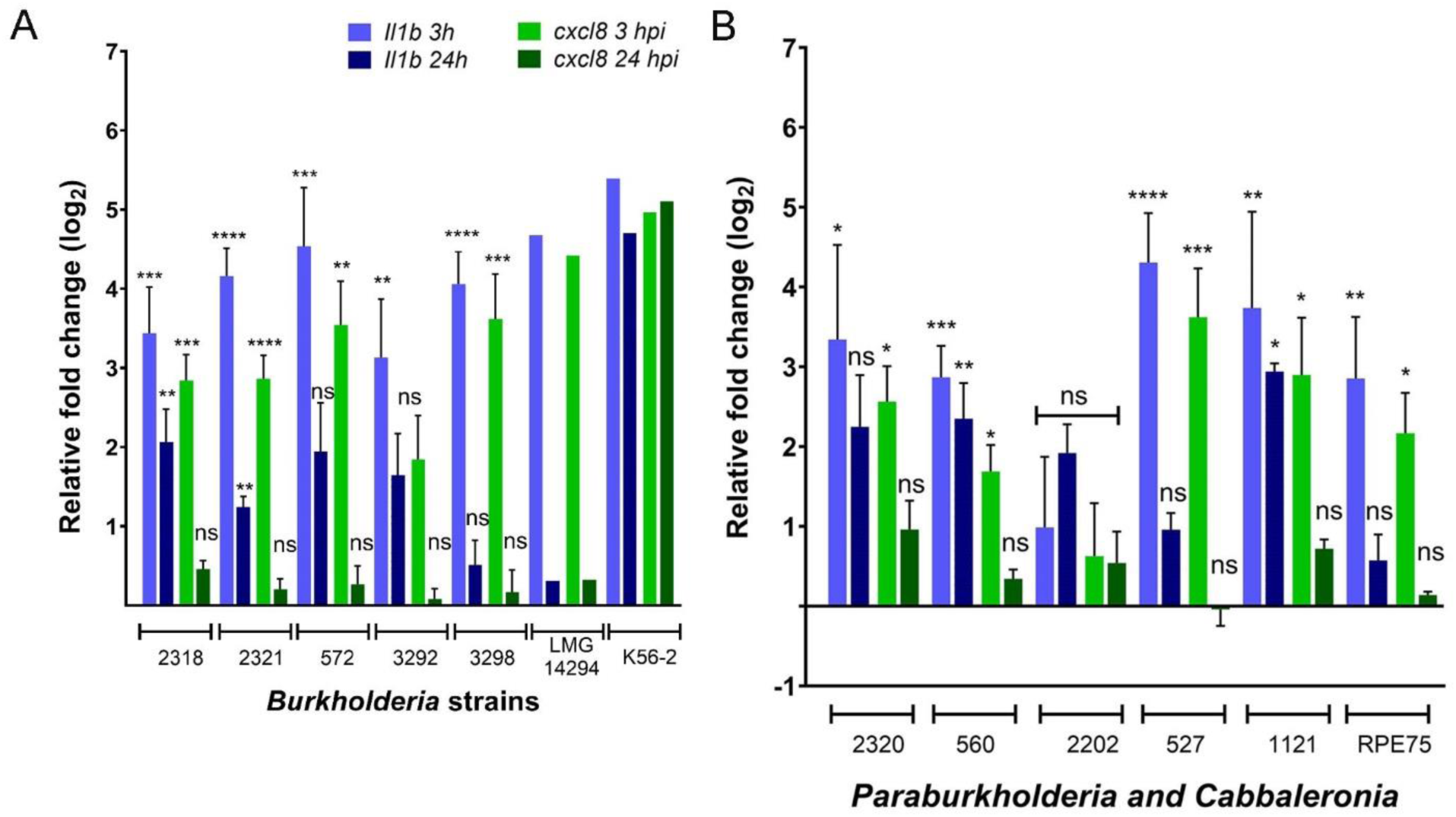
Mean relative cytokine expression levels (in log_2_ fold change) in embryos injected at 3 and 24hpi with the panel of strains. **A.** *Burkholderia*, **B.** *Paraburkholderia* and *Caballeronia* strains. The average inoculum is the same as those indicated in the legend to Figure 2. For *B. stabilis* LMG14294 and *B. cenocepacia* K56-2 the inoculum was K56-2: 438, LMG14294: 398. *Il1b* (Blue bars) and *cxcl8* (green bars) are normalized to the PBS-injected control group for each strain using One-way Anova with Sidak’s with Multiple Comparisons test. Error bars represent mean with SEM of 3 biological replicates (1 experiment for K56-2 and LMG14294), with triplicates in each experiment. Asterisks above each bar indicate significance compared to the PBS control for each strain at 3 and 24 hpi, respectively. * p≤ 0.1; ** p ≤ 0.01; *** p ≤ 0.001; **** p ≤ 0.0001; ns: not significant.

### Neutrophil quantification

To obtain more information about neutrophil responses, we used Tg(*mpx:eGFP*) transgenic embryos expressing GFP in neutrophils. Neutrophil numbers were quantified in embryos injected with 2 *Paraburkholderia* strains, *P. tropica-*like ABIP 1121 and *P. phytofirmans* ABIP 527, and *B. cepacia* ABIP 2318. Images were obtained at 1, 2, 3, and 4 dpi using fluorescence microscopy (**Figure 6**). Neutrophils were quantified using Imaris (see Materials and methods, **Figure 7**). Injection of all three strains resulted in a significantly increased number of neutrophils compared to the non-infected control embryos.

**Figure 6.**
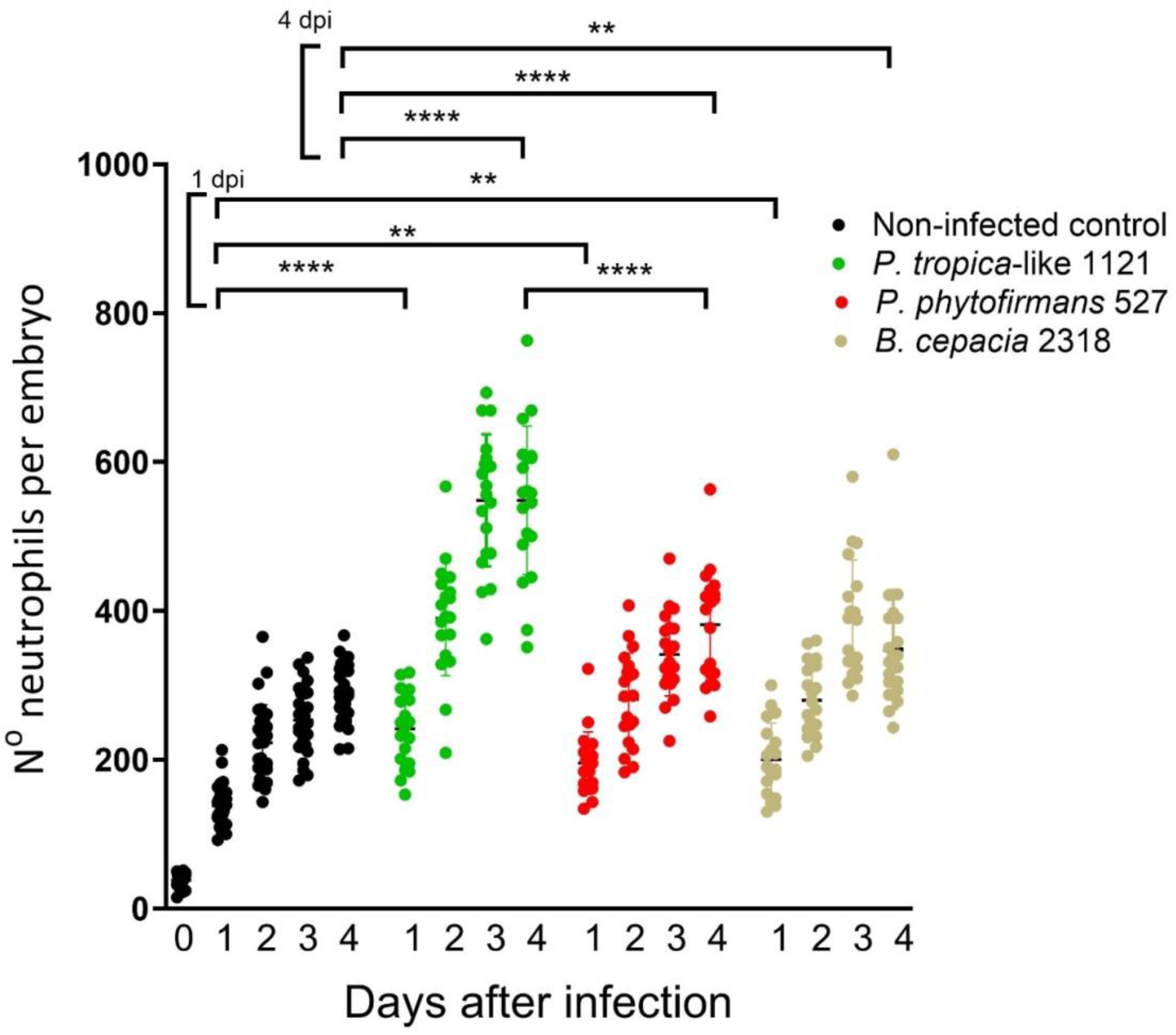
Neutrophil quantification in *mpx-GFP* embryos injected with *P. tropica*-like ABIP 1121, *P. phytofirmans* ABIP 527 and *B. cepacia* ABIP 2318, as well as non-infected control embryos. One-way ANOVA comparing neutrophil numbers at 1- and 4-days post infection. ** p ≤ 0.01; *** p ≤ 0.001; **** p ≤ 0.0001.

**Figure 7.**
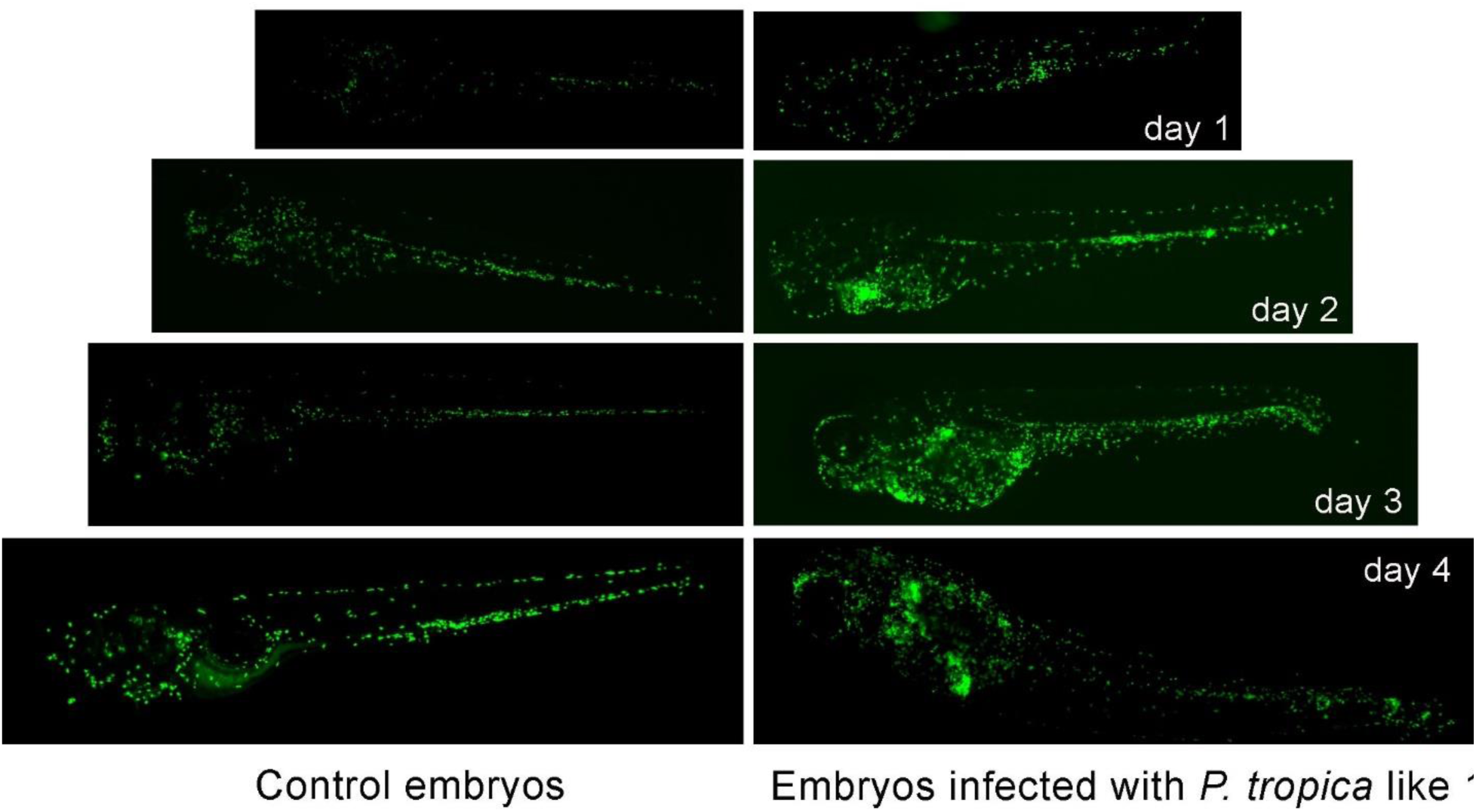
Visualisation of neutrophils during infection. Representative Tg(*mpg-GFP*) embryos, control (left) and infected with *B. tropica*-like ABIP 1121 (right) at 1, 2, 3, and 4 dpi. Control embryos match the same age as infected embryos.

Embryos infected with *P. tropica*-like ABIP 1121 showed the highest increase in neutrophil numbers compared to non-infected embryos at all analysed time points (**Figure 6 and 7**), and with significantly more neutrophils at 4 dpi than *P. phytofirmans* ABIP 527 and *B. cepacia* ABIP 2318. As can be seen in Figure 7, besides the high number of neutrophils in *P. tropica*-like infected embryos compared to non-infected controls embryos, imaging of infected embryos showed neutrophil recruitment at infection sites and sites of abscess formation, in agreement with sustained inflammation.

## Discussion

The aim of this study was to analyse the behavior and potential virulence of a panel of environmental *Burkholderia* s.l. strains (5 *Burkholderia* s.s., 5 *Paraburkholderia*, and a *Caballeronia* strain) in the vertebrate zebrafish embryo model. Our previous work has shown that zebrafish embryos are an excellent model to study virulence and host interactions with *Burkholderia* s.s. strains ^34,44,49^. These earlier studies showed that different clinical and environmental Bcc isolates gave a wide spectrum of infection profiles which ranged from highly virulent, killing embryos in a few days (clinical isolates belonging to the epidemic ET12 lineage, including *B. cenocepacia* K56-2) to persistent non-fatal infections with low pro-inflammatory responses (including the clinical isolates *B. stabilis* LMG14294 and *B. vietnamiensis* FC144)^34,44^. Two environmental isolates *B. cenocepacia* MCO-3 and *B. lata* 383 had intermediate phenotypes, causing 40% and 60 % embryo mortality, respectively ^61^.

Our study revealed 3 strains (*B. cepacia*, *P. caribensis,* and *C. insecticola*) which were cleared from the infected embryos within 3 days, although *C. insecticola* was occasionally found at later stages in the host as micro colonies, possibly in macrophages. *P. tropica-*like was particularly virulent. Although *P. tropica-* like ABIP 1121 did not cause the same degree of mortality as some of the Bcc strains described in our previous studies (K56-2, MCO-3, *B. lata*)^34,61^, it elicited strong inflammatory responses and caused abscess formation, with low vital prognosis for most infected embryos at the end of the experiment. Also, intravenous injection with *P. phytofirmans* ABIP 527 resulted in localized infection sites with evidence of bacterial replication and abscess formation but with lower incidence than *P. tropica*-like ABIP 1121. The remaining 6 isolates were able to survive and persist in the presence of the innate immune system of the young zebrafish, with minimal changes in pro-inflammatory cytokine gene expression at 1 dpi, no visual tissue damage, and no or low mortality, very similar to what has been shown for *B. stabilis* LMG14294 (^34^ and this study). Bacteria from these strains were observed as single or multiple micro-colonies, suggesting that some of the injected bacteria were able to avoid degradation by host immune cells and establish a replication niche, while occasionally more heavily infected embryos were observed. We earlier established that, after 8 dpi, embryos chronically infected with *B. stabilis* LMG14294 showed exacerbation of the infection in about 10% of infected embryos^34^; it would be interesting to study the persistent character of the panel of strains over longer time periods to get more insight into chronicity and exacerbation of infection of the different *Burkholderia* s.l. strains.

All strains evoked a rapid increase in expression of pro-inflammatory cytokine genes *il1b* and *cxcl8* at 3 hpi, as expected after inoculation. Although not significant, Bcc strains seemed to elicit a slightly stronger increase at this early time point than *Paraburkholderia* strains, hinting at the presence of bacterial factors that elicit stronger host responses towards Bcc. *P. tropica*-like ABIP1121, showed the highest levels of *il1b* expression at 24 hpi, in agreement with the observed virulence of this strain. It has been shown previously that acute infection by *B. cenocepacia* K56-2 is associated with systemic phagocyte death within 24 hpi, whereas persistent infection with *B. stabilis* LMG14294 is correlated with neutrophilic inflammation ^44^. In this study, three analysed strains, including *B. cepacia* ABIP 2318, which was cleared by day 3 from the embryos, showed inflammation with significant increase in neutrophil numbers in response to the infection. *P. tropica*-like elicited the highest increase in neutrophils, in agreement with the sustained expression of *il1b*, and higher than that reported earlier by *B. stabilis* LMG14294 ^44^; however the infection did not result in phagocyte death as seen during acute infections caused by K56-2. The reason for the K56-2-induced phagocyte death is however not yet clear but could be due to excessive inflammatory responses and immune cell apoptosis, or caused by bacterial secretion of hemolytic factors.

With classically extracellular bacterial species such as *Pseudomonas aeruginosa* and *Staphylococcus aureus* infectious doses of >1000 CFU are generally used for blood stream infections in zebrafish embryos ^62,63^. For *Burkholderia*, we have previously shown that 50-200 CFU reproducibly leads to fatal infection with virulent strains such as the clinical isolate *B. cenocepacia* K56-2, while other clinical isolates (*B. stabilis* LMG14294, *B. vietnamiensis* FC441) caused infection with constant bacterial burden and no host death in the time course of the experiment. Our earlier results also demonstrated that bacteria were rapidly phagocytosed by macrophages ^59^ and that *B. cenocepacia* required an intramacrophage replication stage to develop acute fatal infection in zebrafish embryos ^44^. To allow injected bacteria to be phagocytosed by the available macrophages after intravenous injection, the infection experiments in this study were performed with a relatively low infection dose of 200-400 CFU. However, injection of a higher dose would most likely aggravate the infection, as hinted by the occasional death of embryos with 600 CFU *B. stabilis* LMG14294 in this study (Figure 1A), not observed with an inoculum of 50-200 CFU^32^.

It will be difficult to extrapolate with accuracy whether *Paraburkholderia* strains might be pathogenic to humans based on experiments using a single animal model, whether it is a non-vertebrate, zebrafish or mammalian model. As example, *B. multivorans* and *B. cenocepacia*, the two most prevalent species infecting CF patients, have shown different outcomes for different isolates in various infection models. Both clinical and environmental isolates of *B. multivorans* can survive and replicate in murine macrophages, similar to *B. cenocepacia* epidemic isolates, including K56-2 ^64^. Another *B. multivorans* CF isolate has been shown to better persist in an intranasal infection model in healthy BALB/c mice than *B. cenocepacia* C6433, a CF isolate that was cleared in the mouse lung ^65^. However, *B. multivorans* was cleared in a mouse model of chronic granulomatous disease, in contrast to *B. cenocepacia* K56-2 ^36^, which has been shown to be virulent in multiple models. In *G. mellonella* and *C. elegans*, *B. multivorans* strains were shown to be relatively avirulent compared to *B. cenocepacia* strains^31,32^. The two strains closest to *B. multivorans* used in this study persisted at different levels, but were not virulent as any of the previously analysed *B. cenocepacia* strains in the zebrafish infection model. Given the previous comments, this does not guarantee that they would be non-pathogenic for humans.

Unexpectedly, amongst our strain panel, the most virulent infection with low mortality rate and poor prognosis for the rest of the infected embryos was caused by a *Paraburkholderia* strain. Recently, Wallner et al^53^ analyzed the transcriptome response of 6 *Burkholderia* s.s. and 6 *Paraburkholderia* strains in their response to root exudates of their hosts. They found no specific gene expression signature for either the *Burkholderia s.s.* or *Paraburkholderia* genus, but rather strain-specific responses to root exudates with few commonly regulated functions ^54^. Interestingly, they found that *P. tropica*-like ABIP659 showed the strongest response to root exudates in terms of numbers of differentially expressed genes (up- and down-regulated), showing its ability to adapt during interaction with plants. Together with its infection phenotype in zebrafish, *P. tropica*-like shows potential to establish host-pathogen interactions in both plant and animal models, though it has no pathogenic behaviour in rice ^54^. In addition, based on the isolation of its close relative *P. tropica* from human infection ^14^ that has plant-growth promoting activity for barley ^13^, the observed virulence for *P. tropica*-like in the zebrafish might be a strong candidate for prediction of virulence in humans. We also observed that *B. phytofirmans* PsJN can cause infection with occasional abscess formation in the zebra fish model. While this strain has been proposed as a bio-fungicide for the control of gray mold disease in grapevine^11^, our data suggest further risk analysis is needed for the safe introduction of such strains in the environment.

The ability of the *Burkholderia* s.l. in general to establish symbiotic interactions with the slime mould *Dictyostelium discoideum* may predispose these bacteria to survive in the hostile intracellular macrophage environment ^66^, as shown here for *P. tropica*-like, which is able to survive and replicate in macrophages. Here, amongst a randomly selected panel of 5 *Burkholderia* s.s. and 5 *Paraburkholderia* isolates, we found no specific differences in bacterial persistence following phylogenetic classifications, suggesting the *Burkholderia* s.l. are generally resistant to degradation by phagocytes. The zebrafish embryo model is particularly interesting to study interactions with macrophages and neutrophils in the context of an innate immune response, and will provide information of the ability of bacteria to resist phagocytic killing in an in vivo model. Studies are also required to determine the bacterium’s ability to survive in the context of an adaptive immune response. More detailed analysis of potential virulence factors, genomic comparisons and analysis of a much larger panel of strains in different animal models will be required to further explore pathogenic potential within the different genera belonging to the *Burkholderia* s.l.. It should, however be remembered that a characteristic thought to be ‘beneficial’ in one condition may be a virulence factor in another. This is highlighted by the antifungal activity cluster AFC, encoded by multiple Bcc species ^60^, which is essential for virulence in rats, plants and insects ^67,68^, and using zebrafish embryos, AFC was shown to be essential for the transition from persistent to acute fatal infection ^60^. It can currently not be excluded that other *Paraburkholderia* strains are potentially pathogenic for humans, and thus phylogenetic classifications cannot be judged as a safety border for pathogenic infections, supporting the opinion of Eberl and Vandamme that “independent of a strain’s phylogenetic status thorough characterization of a strain will be required before it can be considered safe” ^8^.

## Author Contributions.

Conceptualization: A.V., L.M.

Sample collection: L.M

Methodology: J.G., J.B., A.V.

Data analysis: J.G., A.V.

Writing—draft: J.G., A.V.

Writing—review and editing: A.V., L.M.

Funding acquisition: A.V. L.M.

All authors have read and agreed to the published version of the manuscript.

## Acknowledgements

The authors thank Adrian Wallner and Agnieszka Klonowska for providing some of the DSRed derivative strains. The strains were obtained from the IRD and follows the guidelines of the Nagoya protocol regarding access and benefit-sharing. The laboratories of AV and LM were financed by the Agence Nationale de Recherche (ANR, BURKADAPTproject - ANR-19-CE20-0007), from which JG received a PhD grant. Nimes Metropole, UM and INSERM contributed to financing the zebrafish facility. We thank David O’Callaghan for critically reviewing the manuscript.

## References

1. Burkholder, W.H. (1950). Sour skin, a bacterial rot of onion bulbs. Phytopathology 40, 115–118.

2. Sawana, A., Adeolu, M., and Gupta, R.S. (2014). Molecular signatures and phylogenomic analysis of the genus *burkholderia*: Proposal for division of this genus into the emended genus *burkholderia* containing pathogenic organisms and a new genus *paraburkholderia* gen. nov. harboring environmental species. Front. Genet. 5, 1–22.

3. Dobritsa, A.P., and Samadpour, M. (2016). Transfer of eleven species of the genus *Burkholderia* to the genus *Paraburkholderia* and proposal of *Caballeronia* gen. nov. to accommodate twelve species of the genera *Burkholderia* and *Paraburkholderia*. Int. J. Syst. Evol. Microbiol. 66, 2836– 2846.

4. Lopes-Santos, L., Castro, D.B.A., Ferreira-Tonin, M., Corrêa, D.B.A., Weir, B.S., Park, D., Ottoboni, L.M.M., Neto, J.R., and Destéfano, S.A.L. (2017). Reassessment of the taxonomic position of *Burkholderia andropogonis* and description of *Robbsia andropogonis* gen. nov., comb. nov. Antonie van Leeuwenhoek, Int. J. Gen. Mol. Microbiol. 110, 727–736.

5. Estrada-de los Santos, P., Palmer, M., Chávez-Ramírez, B., Beukes, C., Steenkamp, E.T., Briscoe, L., Khan, N., Maluk, M., Lafos, M., Humm, E., et al. (2018). Whole genome analyses suggests that *Burkholderia* sensu lato contains two additional novel genera (*Mycetohabitans* gen. nov., and *Trinickia* gen. nov.): Implications for the evolution of diazotrophy and nodulation in the *Burkholderiaceae*. Genes. 9 (8): 389.

6. Lin, Q.H., Lv, Y.Y., Gao, Z.H., and Qiu, L.H. (2020). *Pararobbsia silviterrae* gen. nov., sp. nov., isolated from forest soil and reclassification of *Burkholderia alpina* as *Pararobbsia alpina* comb. nov. Int. J. Syst. Evol. Microbiol. 70, 1412–1420.

7. Mullins, A.J., and Mahenthiralingam, E. (2021). The Hidden Genomic Diversity, Specialized Metabolite Capacity, and Revised Taxonomy of *Burkholderia* Sensu Lato. Front. Microbiol. 12. 726847.

8. Eberl, L., and Vandamme, P. (2016). Members of the genus *Burkholderia*: good and bad guys. F1000 Research 5, 1007.

9. De Lajudie, P., and Young, J. (2017). International committee on systematics of prokaryotes subcommittee for the taxonomy of *Rhizobium* and *Agrobacterium* minutes of the meeting, Budapest, 25 August 2016. Int. J. Syst. Evol. Microbiol. 67, 2485–2494.

10. Angus, A.A., Agapakis, C.M., Fong, S., Yerrapragada, S., Estrada-de Los Santos, P., Yang, P., Song, N., Kano, S., Caballero-Mellado, J., De Faria, S.M., et al. (2014). Plant-associated symbiotic *Burkholderia* species lack hallmark strategies required in mammalian pathogenesis. PLoS One 9(1):e83779

11. Vilanova, L.C.M., Rondeau, M., Robineau, M., Guise, J.F., Lavire, C., Vial, L., Fontaine, F., Clément, C., Jacquard, C., Esmaeel, Q., et al. (2022). *Paraburkholderia phytofirmans* PsJN delays *Botrytis cinerea* development on grapevine inflorescences. Front. Microbiol 13, 1030982.

12 Kaur, C., Selvakumar, G., and Ganeshamurthy, A. (2017). Burkholderia to Paraburkholderia: The Journey of a Plant-Beneficial-Environmental Bacterium. In Recent advances in Applied Microbiology, pp. 213–228.

13. García, S.S., Bernabeu, P.R., Vio, S.A., Cattelan, N., García, J.E., Puente, M.L., Galar, M.L., Prieto, C.I., and Luna, M.F. (2020). *Paraburkholderia tropica* as a plant-growth–promoting bacterium in barley: characterization of tissues colonization by culture-dependent and -independent techniques for use as an agronomic bioinput. Plant Soil 451, 89–106.

14. Deris, Z.Z., Van Rostenberghe, H., Habsah, H., Noraida, R., Tan, G.C., Chan, Y.Y., Rosliza, A.R., and Ravichandran, M. (2010). First isolation of *Burkholderia tropica* from a neonatal patient successfully treated with imipenem. Int. J. Infect. Dis. 14, 73–74.

15. Coenye, T., Laevens, S., Willems, A., Ohlén, M., Hannant, W., Govan, J.R.W., Gillis, M., Falsen, E., and Vandamme, P. (2001). *Burkholderia fungorum* sp. nov. and *Burkholderia caledonica* sp. nov., two new species isolated from the environment, animals and human clinical samples. Int. J. Syst. Evol. Microbiol. 51, 1099–1107.

16. Gerrits, G.P., Klaassen, C., Coenye, T., Vandamme, P., and Meis, J.F. (2005). *Burkholderia fungorum* septicemia. Emerg. Infect. Dis. 11, 1115–1117.

17. Peeters, C., Meier-Kolthoff, J.P., Verheyde, B., De Brandt, E., Cooper, V.S., and Vandamme, P. (2016). Phylogenomic study of *Burkholderia glathei*-like organisms, proposal of 13 novel *Burkholderia* species and emended descriptions of B*urkholderia sordidicola, Burkholderia zhejiangensis, and Burkholderia grimmiae*. Front. Microbiol. 7, 1–19.

18. Eissa, M. (2024). Genus *Burkholderia*: A Double-Edged Sword with Widespread Implications for Human Health, Agriculture, and the Environment. J. Biol. Res. Rev. 1, 79–96.

19. Mahenthiralingam, E., Baldwin, A., and Dowson, C.G. (2008). *Burkholderia cepacia* complex bacteria: Opportunistic pathogens with important natural biology. J. Appl. Microbiol. 104, 1539– 1551.

20. Scoffone, V.C., Chiarelli, L.R., Trespidi, G., Mentasti, M., Riccardi, G., and Buroni, S. (2017). *Burkholderia cenocepacia* Infections in Cystic Fibrosis Patients : Drug Resistance and Therapeutic Approaches. Front. Microbiol. 8, 1–13.

21. Blanchard, A.C., Tang, L., Tadros, M., Muller, M., Spilker, T., Waters, V.J., Lipuma, J.J., and Tullis, E. (2020). *Burkholderia cenocepacia* ET12 transmission in adults with cystic fibrosis. Thorax 75, 88– 90.

22. Greenberg, D.E., Goldberg, J.B., Stock, F., Murray, P.R., Holland, S.M., and Lipuma, J.J. (2009). Recurrent *Burkholderia* infection in patients with chronic granulomatous disease: 11-Year experience at a large referral center. Clin. Infect. Dis. 48, 1577–1579.

23. Tavares, M., Kozak, M., Balola, A., and Sá-Correia, I. (2020). *Burkholderia cepacia* complex bacteria: A feared contamination risk in water-based pharmaceutical products. Clin. Microbiol. Rev. 33, e00139–19.

24. Ali, M. (2016). *Burkholderia cepacia* in Pharmaceutical Industries. Int. J. Vaccines Vaccin. 3, 00064.

25. Karanth, S.S., Regunath, H., Chawla, K., and Prabhu, M. (2012). A rare case of community acquired *Burkholderia cepacia* infection presenting as pyopneumothorax in an immunocompetent individual. Asian Pac. J. Trop. Biomed. 2, 166–168.

26. Hammoud, M., Fares, Y., Atoui, R., and Dabboucy, B. (2019). *Burkholderia cepacia* as a cause of pyogenic spondylodiscitis in immunocompetent patients: a single-institution case series and literature review. J. Spine Surg. 5, 372–377.

27. Ranjan, R., Chowdhary, P., and Kamra, A. (2017). Community acquired *Burkholderia cepacia* bacteraemia presenting as MODS in an immunocompetent individual: An unusual case. J. Clin. Diagnostic Res. 11, DD01–DD02.

28. Valvano, M.A. (2015). Intracellular survival of *Burkholderia cepacia* complex in phagocytic cells. Can. J. Microbiol. 615, 607–615.

29. Castonguay-Vanier, J., Vial, L., Tremblay, J., and Déziel, E. (2010). *Drosophila melanogaster* as a model host for the *Burkholderia cepacia* complex. PLoS One 5. e11467.

30. Uehlinger, S., Schwager, S., Bernier, S.P., Riedel, K., Nguyen, D.T., Sokol, P.A., and Eberl, L. (2009). Identification of specific and universal virulence factors in *Burkholderia cenocepacia* strains by using multiple infection hosts. Infect. Immun. 77, 4102–4110.

31. Seed, K.D., and Dennis, J.J. (2008). Development of *Galleria mellonella* as an alternative infection model for the *Burkholderia cepacia* complex. Infect. Immun. 76, 1267–1275.

32. Cardona, S.T., Wopperer, J., Eberl, L., and Valvano, M.A. (2005). Diverse pathogenicity of *Burkholderia cepacia* complex strains in the *Caenorhabditis elegans* host model. FEMS Microbiol. Lett. 250, 97–104.

33. Bragonzi, A. (2010). Murine models of acute and chronic lung infection with cystic fibrosis pathogens. Int. J. Med. Microbiol. 300, 584–593.

34. Vergunst, A.C., Meijer, A.H., Renshaw, S.A., and O’Callaghan, D. (2010). *Burkholderia cenocepacia* creates an intramacrophage replication niche in zebrafish embryos, followed by bacterial dissemination and establishment of systemic infection. Infect. Immun. 78, 1495–1508.

35. Kothe, M., Antl, M., Huber, B., Stoecker, K., Ebrecht, D., Steinmetz, I., and Eberl, L. (2003). Killing of *Caenorhabditis elegans* by *Burkholderia cepacia* is controlled by the *cep* quorum-sensing system. Cell. Microbiol. 5, 343–351.

36. Sousa, S.A., Ulrich, M., Bragonzi, A., Burke, M., Worlitzsch, D., Leitão, J.H., Meisner, C., Eberl, L., Sá-correia, I., and Döring, G. (2007). Virulence of *Burkholderia cepacia* complex strains in *gp91phox-*/-mice. Cell. Microbiol. 9, 2817–2825.

37. Bernier, S.P., Silo-suh, L., Woods, D.E., Ohman, D.E., and Sokol, P. a (2003). Comparative Analysis of Plant and Animal Models for Characterization of *Burkholderia cepacia* Virulence. Society 71, 5306–5313.

38. DuBose, J.G., Robeson, M.S., Hoogshagen, M., Olsen, H., and Haselkorn, T.S. (2022). Complexities of Inferring Symbiont Function: Paraburkholderia Symbiont Dynamics in Social Amoeba Populations and Their Impacts on the Amoeba Microbiota. Appl. Environ. Microbiol. 88, e0128522.

39. Meijer, A.H., and Spaink, H.P. (2011). Host-Pathogen Interactions Made Transparent with the Zebrafish Model. Curr. Drug Targets 12, 1000–1017.

40. Torraca, V., and Mostowy, S. (2018). Zebrafish Infection: From Pathogenesis to Cell Biology. Trends Cell Biol. 28, 143–156.

41. Renshaw, S.A., and Trede, N.S. (2012). A model 450 million years in the making: zebrafish and vertebrate immunity. Dis. Model. Mech. 5, 38–47.

42. Van Der Vaart, M., Spaink, H.P., and Meijer, A.H. (2012). Pathogen recognition and activation of the innate immune response in zebrafish. Adv. Hematol. 2012. 159807.

43. Franza, M., Varricchio, R., Alloisio, G., Simone, G. De, Bella, S. Di, Ascenzi, P., and Masi, A. di (2024). Zebrafish (*Danio rerio*) as a Model System to Investigate the Role of the Innate Immune Response in Human Infectious Diseases. Int. J. Mol. Sci. 25. 12008

44. Mesureur, J., Feliciano, J.R., Wagner, N., Gomes, M.C., Zhang, L., Blanco-Gonzalez, M., Van Der Vaart, M., O’callaghan, D., Meijer, A.H., and Vergunst, A.C. (2017). Macrophages, but not neutrophils, are critical for proliferation of Burkholderia cenocepacia and ensuing host-damaging inflammation. 13(6): e1006437.

45. Schwab, U., Abdullah, L.H., Perlmutt, O.S., Albert, D., William Davis, C., Arnold, R.R., Yankaskas, J.R., Gilligan, P., Neubauer, H., Randell, S.H., et al. (2014). Localization of *Burkholderia cepacia* complex bacteria in cystic fibrosis lungs and interactions with Pseudomonas aeruginosa in hypoxic mucus. Infect. Immun. 82, 4729–4745.

46. Sajjan, U., Corey, M., Humar, A., Tullis, E., Cutz, E., Ackerley, C., and Forstner, J. (2001). Immunolocalisation of *Burkholderia cepacia* in the lungs of cystic fibrosis patients. J. Med. Microbiol. 50, 535–546.

47. R Renshaw, S.A., Loynes, C.A., Trushell, D.M.I., Elworthy, S., Ingham, P.W., and Whyte, M.K.B. (2006). A transgenic zebrafish model of neutrophilic inflammation. Blood 108, 3976–3978.

48. Nguyen-Chi, M., Phan, Q.T., Gonzalez, C., Dubremetz, J.F., Levraud, J.P., and Lutfalla, G. (2014). Transient infection of the zebrafish notochord with *E. coli* induces chronic inflammation. DMM Dis. Model. Mech. 7, 871–882.

49. Mesureur, J., and Vergunst, A.C. (2014). Zebrafish embryos as a model to study bacterial virulence. Methods Mol. Biol. 1197, 41–66.

50. Sondo, M., Wonni, I., Koïta, K., Rimbault, I., Barro, M., Tollenaere, C., Moulin, L., and Klonowska, A. (2023). Diversity and plant growth promoting ability of rice root-associated bacteria in Burkina-Faso and cross-comparison with metabarcoding data. PLoS One 18, e0287084.

51. Viallard, V., Poirier, I., Cournoyer, B., Haurat, J., Wiebkin, S., Ophel-keller, K., and Balandreaul, J. (1998). *Burkholderia graminis* sp. nov., a rhizospheric *Burkholderia* species, and reassessment of [*Pseudomonas*] *phenazinium*, [*Pseudomonas*] *pyrrocinia* and [*Pseudomonas*] *glathei* as *Burkholderia*. Int J Syst Bacteriol 48, 549–563.

52. Mattos, K.A., Pádua, V.L., Romeiro, A., Hallack, L.F., Neves, B.C., Ulisses, T.M., Barros, C.F., Todeschini, A.R., Previato, J.O., and Mendonça-previato, L. (2008). Endophytic colonization of rice (*Oryza sativa* L.) by the diazotrophic bacterium *Burkholderia kururiensis* and its ability to enhance plant growth. Ann. Brazilian Acad. Sci. 80, 477–493.

53. King, E., Wallner, A., Guigard, L., Rimbault, I., Parrinello, H., Klonowska, A., Moulin, L., and Czernic, P. (2023). *Paraburkholderia phytofirmans* PsJN colonization of rice endosphere triggers an atypical transcriptomic response compared to rice native *Burkholderia* s.l. endophytes. Sci. Rep. 13, 1–17.

54. Wallner, A., Klonowska, A., Guigard, L., King, E., Rimbault, I., Ngonkeu, E., Nguyen, P., Béna, G., and Moulin, L. (2023). Comparative genomics and transcriptomic response to root exudates of six rice root-associated *Burkholderia* sensu lato species. Peer Community J. 3. e25.

55. Shibata, T.F., Maeda, T., Nikoh, N., Yamaguchi, K., Oshima, K., Hattori, M., and Nishiyama, T. (2013). Complete Genome Sequence of *Burkholderia* sp. Strain RPE64, bacterial Symbiont of the Bean Bug *Riptortus pedestris*. genomeAnounc 1(4): e00441–13.

56. Wallner, A., King, E., Ngonkeu, E.L.M., Moulin, L., and Béna, G. (2019). Genomic analyses of *Burkholderia cenocepacia* reveal multiple species with differential host-Adaptation to plants and humans. BMC Genomics 20, 1–15.

57. Gillis, M., Van, T.V.A.N., Bardin, R., Goor, M., Hebbar, P., Willems, A., Segers, P., Kersters, K., Heulin, T., and Fernandez, M.P. (1995). Polyphasic Taxonomy in the Genus *Burkholderia* Leading to an Emended Description of the Genus and Proposition of *Burkholderia vietnamiensis* sp. nov. for N,- Fixing Isolates from Rice in Vietnam. Int. J. Syst. bacteriol. 45, 274–289.

58. Darling, P., Chan, M., Cox, A.D., and Sokol, P.A. (1998). Siderophore production by cystic fibrosis isolates of *Burkholderia cepacia*. Infect. Immun. 66, 874–877.

59. Revets, H., Vandamme, P., Van Zeebroeck, A., De Boeck, K., Struelens, M.J., Verhaegen, J., Ursi, J.P., Verschraegen, G., Franckx, H., Malfroot, A., et al. (1996). *Burkholderia* (*Pseudomonas*) *cepacia* and cystic fibrosis: the epidemiology in Belgium. Acta Clin. Belg. 51, 222–230.

60. Gomes Id, M.C., Tasrini, Y., Subramoni, S., Id, K.A., Feliciano Id, J.R., Id, L.E., Sokol, P., O’callaghan, D., and Vergunst Id, A.C. (2018). The *afc* antifungal activity cluster, which is under tight regulatory control of ShvR, is essential for transition from intracellular persistence of *Burkholderia cenocepacia* to acute pro-inflammatory infection. PLoS Pathog 14(12): e1007473.

61. Agnoli, K., Freitag, R., Gomes, M.C., Jenul, C., Suppiger, A., Mannweiler, O., Frauenknecht, C., Janser, D., Vergunst, A.C., and Eberl, L. (2017). Use of synthetic hybrid strains to determine the role of replicon 3 in virulence of the *Burkholderia cepacia* complex. Appl. Environ. Microbiol. 83, 1–17.

62. Prajsnar, T.K., Hamilton, R., Garcia-Lara, J., Mcvicker, G., Williams, A., Boots, M., Foster, S.J., and Renshaw, S.A. (2012). A privileged intraphagocyte niche is responsible for disseminated infection of *Staphylococcus aureus* in a zebrafish model. Cell. Microbiol. 14, 1600–1619.

63. Clatworthy, A.E., Lee, J.S.W., Leibman, M., Kostun, Z., Davidson, A.J., and Hung, D.T. (2009). *Pseudomonas aeruginosa* infection of zebrafish involves both host and pathogen determinants. Infect. Immun. 77, 1293–1303.

64. Schmerk, C.L., and Valvano, M.A. (2013). *Burkholderia multivorans* survival and trafficking within macrophages. J. Med. Microbiol. 62, 173–184.

65. Chu, K.K., Macdonald, K.L., Davidson, D.J., and Speert, D.P. (2004). Persistence of Burkholderia multivorans within the pulmonary macrophage in the murine lung. 72, 6142–6147.

66. Khojandi, N., Haselkorn, T.S., Eschbach, M.N., Naser, R.A., and Disalvo, S. (2019). Intracellular *Burkholderia* Symbionts induce extracellular secondary infections; driving diverse host outcomes that vary by genotype and environment. ISME J. 13, 2068–2081.

67. Subramoni, S., Nguyen, D.T., and Sokol, P.A. (2011). *Burkholderia cenocepacia* ShvR-regulated genes that influence colony morphology, biofilm formation, and virulence. Infect. Immun. 79, 2984–2997.

68. Thomson, E.L.S., and Dennis, J.J. (2013). Common Duckweed (*Lemna minor*) Is a Versatile High-Throughput Infection Model For the *Burkholderia cepacia* Complex and Other Pathogenic Bacteria. PLoS One 8, e80102.

